# Identification of a Novel PARP14 Site Motif and Glycohydrolase Specificity Using TLC-MALDI-TOF

**DOI:** 10.1101/2023.03.22.533863

**Authors:** Zeeshan Javed, Hannah H. Nguyen, Kiana Harker, Christian M. Mohr, Pia Vano, Sean R. Wallace, Clarissa Silvers, Colin Sim, Soumya Turumella, Ally Flinn, Ian Carter-O’Connell

## Abstract

Transfer of ADP-ribose (ADPr) from nicotinamide adenine dinucleotide (NAD^+^) to target proteins is mediated by a class of human poly-ADP-ribose polymerases, PARPs, and removal of ADPr is catalyzed by a family of glycohydrolases. Although thousands of potential ADPr modification sites have been identified using high-throughput mass-spectrometry, relatively little is known about sequence specificity encoded near the modification site. Herein, we present a matrix-assisted laser desorption/ionization time-of-flight (MALDI-TOF) method that facilitates the discovery and validation of ADPr site motifs. We identify a minimal 5-mer peptide sequence that is sufficient to drive PARP14 specific activity while highlighting the importance of the adjacent residues in PARP14 targeting. We measure the stability of the resultant ester bond and show that non-enzymatic removal is sequence independent and occurs within hours. Finally, we use the ADPr—peptide to highlight differential activities within the glycohydrolase family and their sequence specificities. Our results highlight: 1) the utility of MALDI-TOF in motif discovery and 2) the importance of peptide sequence in governing ADPr transfer and removal.

---

ADP-ribosylation is a ubiquitous post-translational modification (PTM) found in a vast array of species.^1^ Despite being one of the first characterized PTMs,^2^ the biochemical mechanisms governing ADP-ribosylation are still not fully understood. In humans, the transfer of ADP-ribose (ADPr) from nicotinamide adenine dinucleotide (NAD^+^) to the protein target is mediated by a family of seventeen poly-ADP-ribose polymerases (PARPs) (Figure 1a).^3^ The PARP family is further subdivided based on whether a single ADPr unit is transferred (mono-PARPs: 4, 7-8, 10-12, 14-16) or whether the initial ADPr can be elongated with multiple ADPr units (PARPs 1, 2, 3, 5a/b).^4^ Initially discovered as DNA base repair enzymes based on the activity of PARP1 in the nucleus,^5^ the PARP family has since been linked to a growing set of biological pathways and disease states.^6,7^ Removal of ADPr is catalyzed by a separate class of glycohydrolases,^8^ which can be subdivided based on the ability to remove poly-ADP-ribose (PAR; PARG),^9^ mono-ADP-ribose (MAR; *macro*D1, *macro*D2, ARH1),^10,11^ or both (TARG1, ARH3).^11,12^ As with the PARP family, ADPr removers have been implicated in a number of essential biological processes.^8^

**Figure 1.**
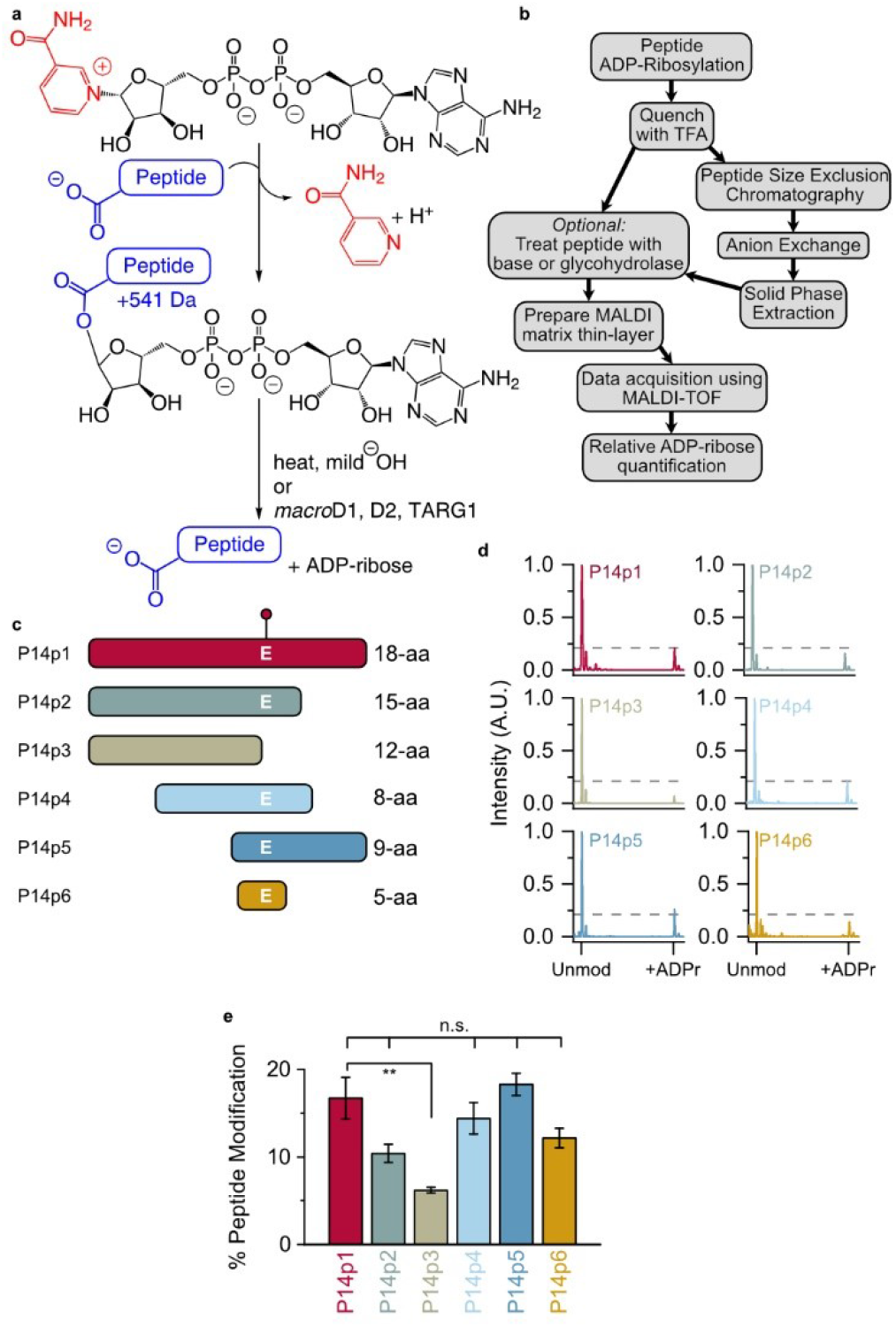
Identification of a minimal P14 selective peptide sequence. (a) ADP-ribosylation (step 1) and hydrolysis reactions (step 2). (b) TLC-MALDI workflow. (c) Peptide truncations used in this study (full sequences in Figure S1). The E acceptor is indicated. (d) P14 and the indicated peptide were incubated in the presence of NAD^+^ and subjected to TLC-MALDI to visualize the resulting increase in *m/z* due to ADPr (+541 Da). The dashed line represents the intensity observed for ADP-ribosylation of P14p1. (e) MS spectra were integrated to determine the relative levels of ADP-ribosylation. The bar graphs depict the fraction of the total peptide that was modified (mean ± S.E.M., *n* = 3). ** represents *p-*value <0.01, two-tailed Student’s *t* test, n.s. represents a non-significant difference.

A fundamental challenge in understanding the function of ADPr signaling in the cell is site identification. As with other types of PTMs, e.g. kinase mediated phosphorylation,^13^ the identification of PARP site motifs would help link distinct modification events to the alteration of target protein function. Recent studies using tandem mass-spectrometry (MS/MS) have approached this challenge by uncovering thousands of potential ADPr sites in the human proteome.^14,15^ Bioinformatic analysis of the resultant data has identified putative site motifs,^16^ but those sites are often weakly enriched and have not been fully validated. The lack of a sequence motif could be due to: (1) a real lack of a consensus target sequence within the PARP family; or (2) the presence of weak motif signatures within an otherwise promiscuous enzyme class; and/or (3) the artificial enrichment of non-physiological sites due to the various interventions required to enrich ADPr sites in MS/MS procedures. There is a difference between enzymes that will modify most of the accessible amino acids on the surface of a target versus ones that modify distinct sites so the presence or absence of site motifs in the PARP family has important implications for PARP function. Additionally, the same factors that dictate ADPr transfer could impact ADPr removal – expanding the importance of sequence motifs more broadly to the glycohydrolases.

Ultra-thin layer chromatography matrix-assisted laser-desorption/ionization time-of-flight (TLC-MADLI) analysis provides a unique opportunity to interrogate ADPr sequence motifs. Unlike MS/MS, the ionization energy employed in MALDI-TOF is not sufficient to disrupt the glycosidic bond, allowing for quantitative assessment of ADPr levels.^17^ The pre-application of a thin-layer of matrix on the steel objective enhances crystal formation in the presence of common contaminants and improves the resultant signal intensity and resolution.^18–20^ Further, the ability to rapidly, and inexpensively, probe peptide substrates in isolation facilitates characterization of each amino acid’s contribution to PARP activity without the confounding input from multiple ADPr sites found on whole proteins. Each of these features of TLC-MALDI is vital for uncovering ADPr site motifs.

We previously used TLC-MALDI to identify an 18-mer peptide (P14p1) that was selectively labeled by PARP14 (P14).^21^ Herein we describe the expansion of our efforts to quantitatively assess the sequence motif preferences of P14 for this peptide (Figure 1b). We demonstrate that a truncated 5 amino acid sequence is sufficient to drive specific modification by P14 and we identify the positions within that sequence that are required for maximal ADPr transfer. We develop a three-step purification strategy to chemoenzymatically synthesize homogenous ADP-ribosylated peptides. We use the resultant ADPr-peptides to assess ADPr removal and describe the effects of specific sequences on both enzymatic and non-enzymatic hydrolysis. We observe that non-enzymatic hydrolysis of ADPr occurs within hours in mild conditions; a result which highlights the reversibility of ADPr *in vivo* and impacts future site identification methods. Taken together, our findings elucidate a novel sequence motif for P14 and demonstrate the wider importance of proximal sequences in ADPr-dependent signaling.

## EXPERIMENTAL SECTION

### ADP-Ribosylation Assay

P14 or P15 (5 µM) was incubated with 500 µM NAD^+^ (Sigma-Aldrich) and the indicated peptide (10 µM) for 10 min at 30 °C in a 16 µL reaction volume consisting of 25 mM HEPES, pH 7.5, 50 mM NaCl, and 0.5 mM TCEP. Reactions were quenched by the addition of 16 µL of 0.1% TFA. 2 µL of sample was mixed with 4 µL of TLC-MALDI buffer (MB, 5 mg/mL α-cyano-4-hydroxycinnamic acid (CHCA, Sigma-Aldrich), 25% acetonitrile, 0.05% TFA, and 5 mM NH_3_PO_4_) and subjected to TLC-MALDI analysis. Details regarding sample cleanup, ADPr—peptide synthesis, and TLC-MALDI acquisition and analysis are included in the Supporting Information.

### Non-enzymatic Hydrolysis Assay

Synthesized ADPr—P14p6 or –P14p8 was equilibrated to 50 µM in 10 mM potassium phosphate, pH 6.0 and diluted in 50 µM BIS-Tris, pH 6.0, 7.0, or 8.0 to a final concentration of 10 µM. ADPr—peptides (2 µL) were incubated at either 37 °C or 25 °C and samples were quenched at 0, 45, 90, 180, or 300 minutes in 4 µL of MB and subjected to TLC-MALDI analysis. For the 4 °C incubations samples were collected at 0, 24, 48, and 72 hours.

### Glycohydrolase Assay

ADPr—P14p6 or –P14p8 (10 µM) was incubated with either *macro*D1, *macro*D2, or TARG1 (500 nM) for 10 min at 30 °C in 2 µL reaction volume consisting of 25 mM BIS-Tris, pH 7.0, 50 mM NaCl, and 0.5 mM TCEP. Reactions were quenched with 4 uL of MB and subjected to TLC-MALDI analysis.

## RESULTS AND DISCUSSION

Initially, a series of P14p1 truncations were designed (P14p2-6) to identify the necessary and sufficient sequence elements required for P14 specific modification (Figure 1c, full sequences in Table S1). Each of the peptides was incubated in the presence of P14 and NAD^+^ and subjected to TLC-MALDI. All of the resulting spectra display varying levels of modification by ADPr as indicated by the expected mass shift of +541 Da (Figures 1a, d). Integration of the unmodified and modified peaks in the MS spectra was used to quantify the relative levels of ADP-ribosylation for each of the truncated peptides (Figure 1e). The loss of residues C-terminal to the modified glutamate (E) results in a decrease in ADPr transfer, though this effect is not significant until the acceptor residue is lost (P14p3) leaving behind a tyrosine (Y) as a potential acceptor. These data are consistent with recent studies demonstrating P14 activity as a Y modifier.^22^ However, it appears that Y is not the preferred acceptor site within this sequence, as its presence as the only P14 target results in a nearly three-fold decrease in modification (from 16.7% to 6.2% of the total peptide). By contrast, the N-terminal portion of the 18-mer can be truncated to an overlap of 2 amino acids with no apparent loss in activity (compare P14p1 to P14p5). While a minor contribution of the final three C-terminal residues in P14 targeting cannot be ruled out, a truncated 5 amino acid sequence surrounding the acceptor E residue showed no significant loss in P14 activity. Importantly, incubation of any of the tested peptides with PARP15, a closely related P14 ortholog, results in almost no transfer of ADPr (Figure S2). These findings demonstrate that the 5-mer sequence selectively targeted by P14 and will be an optimal minimal fragment to assess the effects of proximal sequence on P14 activity.

Next, we assessed the relative contribution of each non-acceptor amino acid on P14 selection. We systematically replaced each of the residues surrounding the E acceptor with alanine (A) and performed TLC-MALDI with P14 as described above (Figure 2a). Our initial experiments with peptide variants that involved the substitution of a charged residue with alanine resulted in spectra that had severely diminished signal intensities compared to the parent peptide (data not shown).

**Figure 2.**
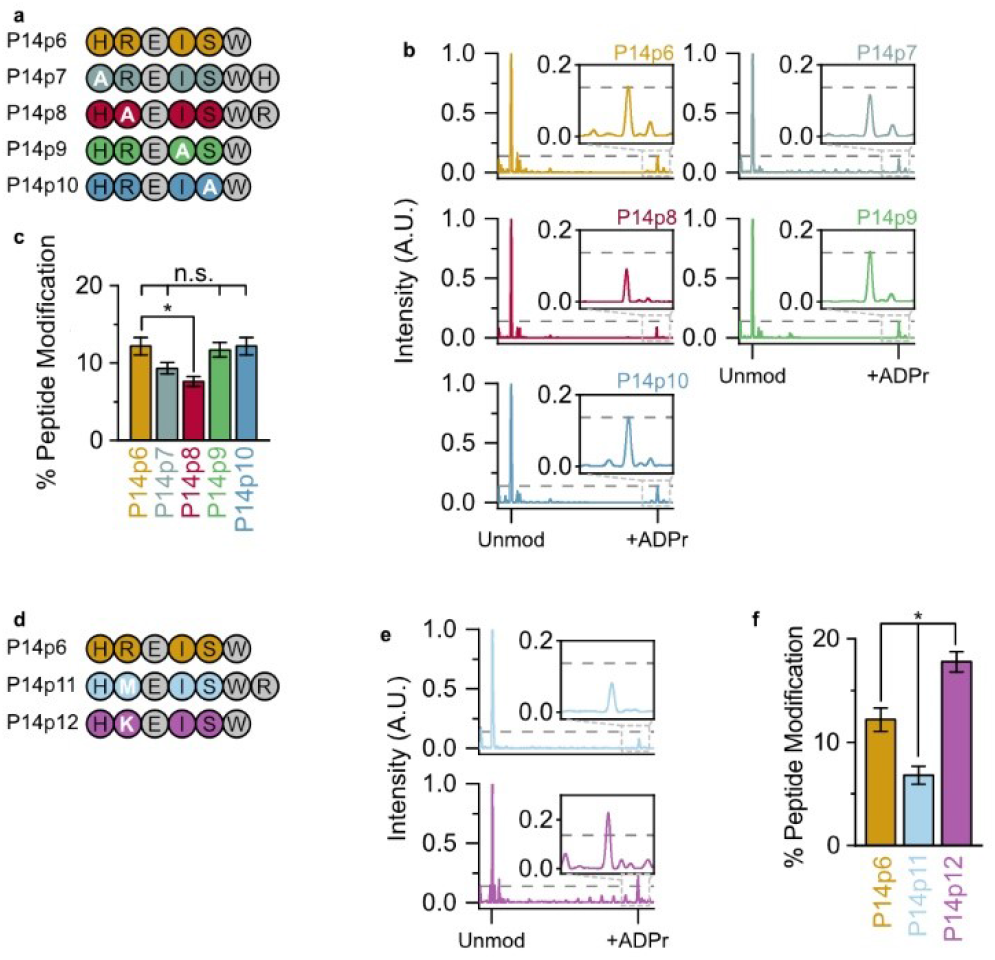
P14 preferentially ADP-ribosylates a basic—K/R—D/E motif. (a) Alanine (A) substituted peptides used in this study. (b) P14 and the indicated peptide were incubated in the presence of NAD^+^ and subjected to TLC-MALDI to visualize the resulting increase in *m/z* due to ADPr (+541 Da). The dashed line represents the intensity observed for ADP-ribosylation of P14p6 and the inset highlights the +ADPr spectra. (c) MS spectra were integrated to determine the relative levels of ADP-ribosylation. The bar graphs depict the fraction of the total peptide that was modified (mean ± S.E.M., *n* = 3). * represents *p-*value <0.05, two-tailed Student’s *t* test, n.s. represents a non-significant difference. (d) Methionine (M) and lysine (K) substituted peptides used in this study. (e) P14p11 and P14p12 MS spectra normalized as in (b). (f) Relative levels of P14p11 and P14 p12 ADP-ribosylation analyzed as in (c).

As ionization in the positive mode used in our MALDI-TOF is dependent on the overall charge of the molecule, we theorized that this lack of signal was caused by the loss of a +1 charge on these peptides. To avoid artifacts due to charge imbalances in the peptides tested, we added back the lost residue to the C-terminus of the peptide while maintaining the alanine substitution (e.g., P14p7 and P14p8). This allowed us to maintain the same charge on each peptide and allowed for direct comparisons in spectral intensity across the screened mutants (Figure 2b). A decrease in ADPr transfer occurred when the acceptor minus two (P14p7) and minus one (P14p8) position was altered to alanine as compared to the native sequence in P14p6 (9.3% or 7.6% peptide modification versus 12.2%, respectively); though the change with P14p7 was not significant (Figure 2c). As both histidine (H) and arginine (R) are positively charged, these data suggest that P14 prefers either a basic and/or larger residue in the two positions N-terminal to the acceptor residue. However, as there was no change in ADPr transfer with an alanine substitution at the two positions C-terminal to the acceptor residue they are non-essential for P14 targeting.

To determine whether the 40% decrease in P14 activity we observed for the P14p8 peptide was due to the positive charge of the arginine (R) or its larger size compared to A, two new peptide substrates were designed with either a methionine (M, P14p11) or lysine (K, P14p12) at the acceptor minus one site (Figure 2d). As with the A substitution, the presence of the larger, though uncharged, M residue results in a 44% decrease in P14 activity. Therefore, the decrease observed with an A substitution was likely not due to the difference in size at this position. The K substitution – which maintains the charge at this position – resulted in a 46% increase in P14 activity. It appears that while R was found at this position in the original sequence, a K residue further enhances activity. Interestingly, the preference for a lysine in the acceptor minus one site has been observed for PARP1^23^ and was one of the putative motifs suggested in prior proteomic analyses of the ADP-ribosylome.^16^ These results demonstrate a general promiscuity in P14 targeting and validate a preference for sequences with a basic–K/R—D/E—X—X motif.

Following the identification of a putative motif for P14-dependent ADPr transfer we determined if this motif influenced enzymatic and non-enzymatic ADPr hydrolysis. Like other ester bonds, the peptide—ADPr linkage is base labile.^24^ However, previous studies on the reversal of the ADPr ester bond routinely used basic conditions well above the pH range observed in the cell (pH > 9),^4,25^ making it difficult to interpret how stable this modification is *in vivo*. The presence of multiple ADPr sites on proteins further complicates the analysis of single-site hydrolysis. We reasoned our TLC-MALDI approach would be well suited to studying the kinetics of single-site ADPr removal without these complications. P14 was used to label both P14p6 and P14p8 homogenously and singly-ADP-ribosylated peptide was purified using a three-step method (see supporting information for details regarding synthesis and purification). The resultant P14p6—ADPr peptide was equilibrated in a range of pH conditions (pH 6 – 8) and the removal kinetics at 37 °C were monitored using TLC-MALDI (Figure 3a). The half-life of ADPr—peptide at pH 8 is 2.2 hours and is only slightly longer at pH 7 (t_1/2_ = 6.0 hours) (Figure 3b). Only by equilibrating the ADPr—peptide in slightly acidic conditions were half-lives in the day range (t_1/2_ = 20.4 hours) observed. The hydrolysis assays were repeated at room temperature (25 °C) to investigate the effect of temperature on the stability of the ADPr—peptide bond. Predictably, lowering the temperature slows the loss of ADPr at all pHs tested, though there is still appreciable removal within mild base (t_1/2_ = 5.6 hours at pH 8) (Figure 3c). However, equilibration in slightly acidic conditions significantly slows the hydrolysis of ADPr and results in a half-life of greater than two days (Figure 3d). Lowering the temperature even further to 4 °C stabilized the modification, though it did not prevent hydrolysis at either pH 8 or 7 (Figure 3e). Of all the conditions tested, only the equilibration of ADPr—P14p6 in acidic conditions on ice seemed to halt non-enzymatic removal (Figure 3f). These results highlight the dynamism of ADP-ribosylation in the cell and have important implications for the methodologies utilized to study ADP-ribosylation (as discussed further below).

**Figure 3.**
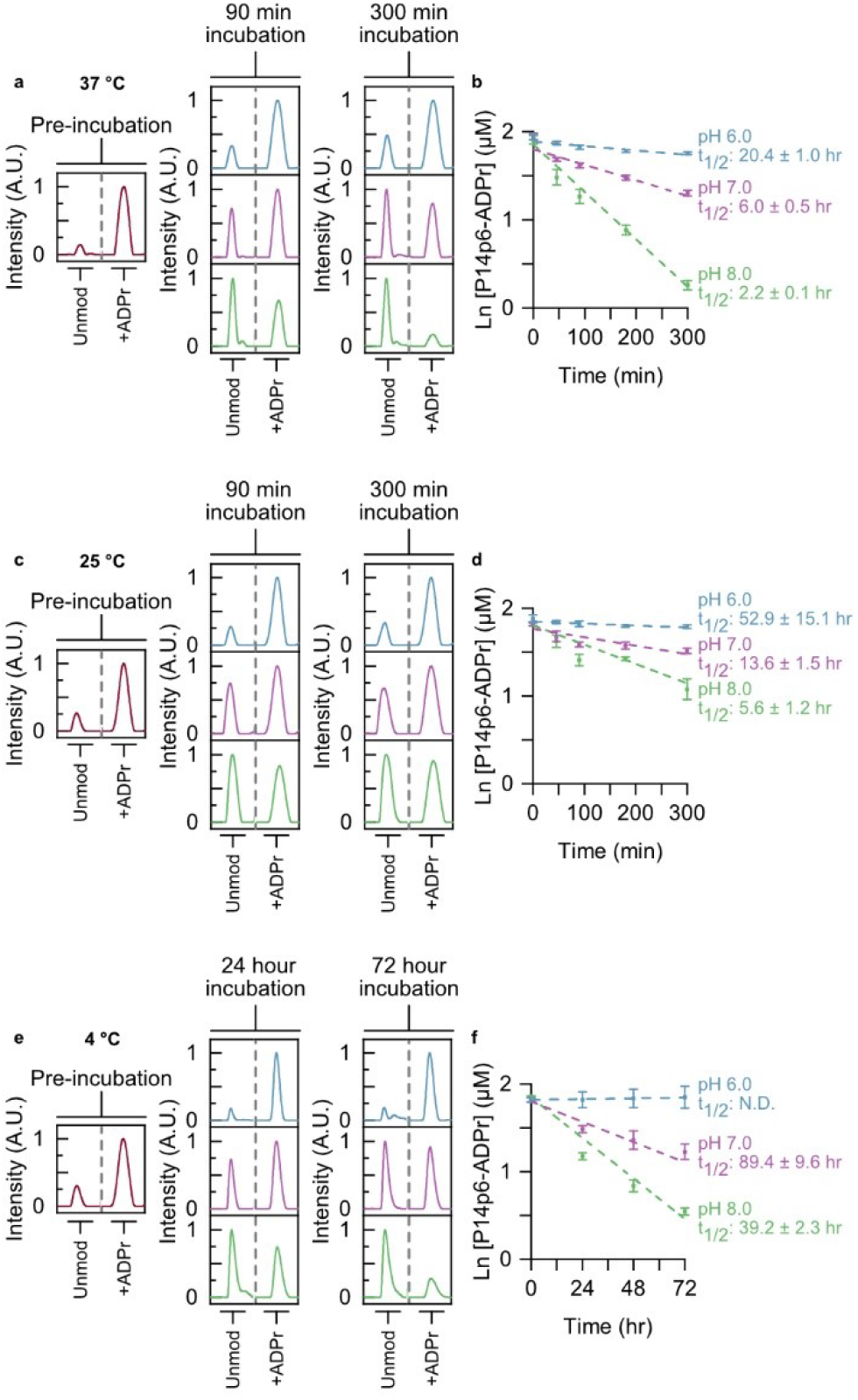
The ADPr—peptide bond is hydrolyzed under mild conditions. (a) Synthesized ADPr—P14p6 peptide was equilibrated at 37 °C and pH 6.0 (blue), pH 7.0 (purple), or pH 8.0 (green) and ADPr hydrolysis was monitored at the indicated times using TLC-MALDI. The unmodified and modified peaks are shown for comparison. (b) MS spectra were integrated to determine the relative levels of ADPr hydrolysis and fit to a pseudo first-order rate expression to determine the half-life of the ADPr modification (mean ± S.E.M., *n* = 3). (c) Incubation at room temperature stabilizes the ADPr—peptide bond. Experiments were performed as in at 25 °C. (d) Determination of ADPr half-lives at 25 °C. (e) Incubation on ice effectively halts ADPr hydrolysis in mild acid. Experiments were performed as in (a) at 4 °C. (f) Determination of ADPr half-lives at 4 °C.

Next, we wanted to determine whether the observed specificity was attributable to either (1) a specific interaction between the acceptor minus one site and P14, or (2) a stabilizing intramolecular interaction between the positive charge on the adjacent R residue and ADPr that slows hydrolysis. To discern this, hydrolysis assays were performed with P14p8 and the rates of hydrolysis for P14p8 were compared to P14p6 (Figure S3). No significant difference was observed in any of the tested conditions between the non-enzymatic rates of hydrolysis for P14p6 and P14p8. As such, the observed increase in ADPr activity seen for P14p6 is likely due to a specific interaction with P14, rather than its susceptibility to hydrolysis.

After surveying non-enzymatic hydrolysis, we moved on to study the effects of proximal sequence on glycohydrolase activity. We limited our study to removal enzymes that had previously been validated as both D/E and mono-ADPr selective: *macro*D1, *macro*D2, TARG1.^10–12^ Each of the glycohydrolases was incubated with either P14p6 or P14p8 and the amount of ADPr that was removed compared to a non-enzymatic control reaction was measured (Figure 4a). All three of the removers showed activity for the P14p6 peptide, with *macro*D2 displaying a 3.8-fold increase in activity compared to *macro*D1 and a 2.0-fold increase compared to TARG1 (Figure 4b). These results suggest a stronger antagonism between P14 and *macro*D2 than the other removers and could indicate a role for *macro*D2 in selective removal of P14 targeted ADPr sites. Then each of the removers was incubated with the P14p8 peptide and ADPr removal was assessed to determine the effects of the acceptor minus one position on glycohydrolase activity. As with P14p6, each of the enzymes tested was active in the presence of the ADP-ribosylated P14p8 peptide (Figure 4a). However, differential sensitivities were observed for the substitution at the acceptor adjacent position, with *macro*D1 displaying no significant difference in activity for P14p6 versus P14p8, while both *macro*D2 and TARG1 had a 30-40% decrease in activity with the altered sequence (Figure 4b). Taken together, this investigation of glycohydrolase activity with P14 specific ADPr—peptides has revealed differences in substrate preference and sensitivity to proximal sequences while demonstrating the utility of TLC-MALDI in elucidating fundamental ADP-ribose biochemistry.

**Figure 4.**
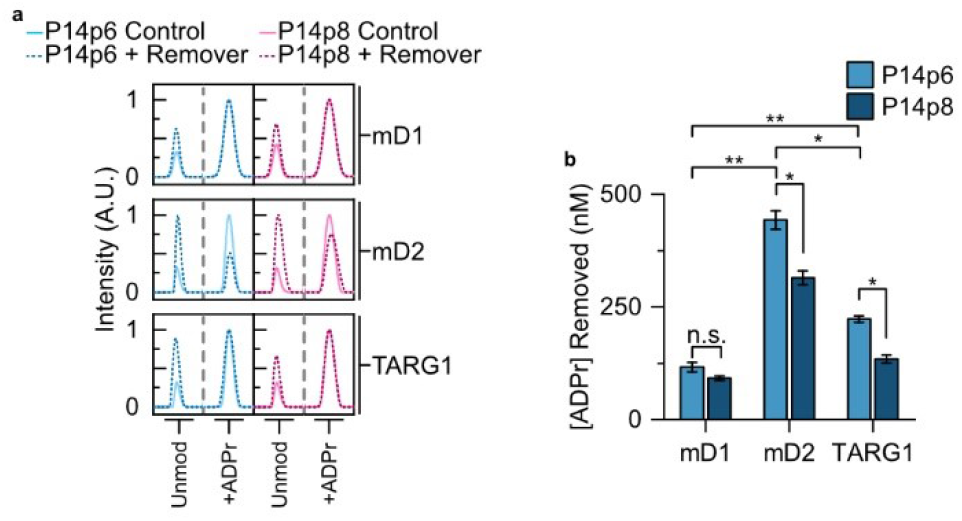
ADPr glycohydrolases display differential preferences for P14 selective sequence motifs. (a) Synthesized ADPr—P14p6 or ADPr—P14p8 peptides (dashed lines) were incubated in the presence of either *macro*D1 (mD1), *macro*D2 (mD2), or TARG1 and subjected to TLC-MALDI to determine the relative levels of hydrolysis. Non-enzymatic controls (solid line) were utilized to ascertain the levels of background hydrolysis. The unmodified and modified peaks are shown for comparison. (b) MS spectra were integrated to determine the relative levels of ADPr hydrolysis. The bar graphs depict the amount of ADPr removal (mean ± S.E.M., *n* = 3). ** represents *p-*value <0.01, two-tailed Student’s *t* test, * represents a *p*-value <0.05, and n.s. represents a non-significant difference.

Herein we demonstrated the applicability of the TLC-MALDI approach in the identification and validation of site motifs within the PARP family. Working with a minimal motif containing a single acceptor site facilitated the determination of the effects of the surrounding amino acids on the activity of P14. While P14 is broadly promiscuous, there is a distinct preference for sites with a basic—K/R—D/E signature. Going forward, it will be interesting to apply this technique to the remaining PARP family members with their own specific target sequences. Completion of this comparative analysis will help reveal the level of specificity within the family and build on the mechanism for PARP targeting in the cell. Further, these results can immediately be used to screen putative sites from proteomic surveys to identify bona fide *in vivo* P14 sites.

One of the main advantages of the described work is the ability to interrogate both attachment and removal of ADPr at a single minimal acceptor site. TLC-MALDI reveals that the esterification of peptide with ADPr results in a far more labile modification than previously described.^4,25^ Understanding how the innate stability of ADPr at D/E residues impacts signal persistence will be of particular interest going forward. Moreover, recent studies have shown that ADPr transfer is not restricted to a single type of amino acid – with PARP activity observed on arginine (R), lysine (K), cysteine (C), histidine (H), serine (S), tyrosine (Y), and the aforementioned D/E.^22,26–30^ Expanding this technique to additional types of ADPr— peptide bond chemistries will help uncover how the unique chemistry at the acceptor site effects PTM lability. Investigation of the interplay between site chemistry and ADPr stability could have important implications for how the kinetics of non-enzymatic removal impact ADPr signal transduction pathways. These results similarly hint at a larger potential pool of underrepresented ADPr sites that have been overlooked due to the instability of the ester bond. When preparing protein libraries for MS/MS analysis it is fairly common to equilibrate the digests in pH neutral conditions at elevated temperatures for more than a day (e.g. overnight treatment with trypsin). Based on the current findings, it is possible that treatment of peptide libraries in this manner would result in a significant loss of ADPr from acidic acceptor sites. In a complex pool of acceptor site chemistries this could result in the over-enrichment of non-ester linkages and an apparent absence of D/E modifications. Therefore, the exploration of the impact of temperature and pH on MS/MS-based site discovery has the potential to identify an underrepresented population of sites.

Finally, the TLC-MALDI method has been successfully adapted from quantifying ADPr transfer to ADPr removal using several different glycohydrolases. As this class of enzymes were thought to be fairly substrate agnostic it was surprising to find differential substrate preferences.^31^ For example, *macro*D1 has been previously shown to remove ADPr from a range of substrates (e.g. DNA, RNA, and protein) and can function as an O-acetyl-ADPr deacetylase.^32^ The data supports the role of *macro*D1 as a promiscuous removal enzyme, but this lack of specificity contributes to a lower activity for the peptidyl substrates analyzed. Recent work by Žaja and co-workers identified an enrichment of *macro*D2 in neuronal cells.^33^ Coupled with the observation that *macro*D2 is the most active remover from the P14 selective ADPr—peptide this could hint at a potential antagonistic role for P14 and *macro*D2 in the brain. Combined, these results have expanded the role for TLC-MALDI in analyzing ADP-ribosylation while providing new insights into the mechanism of target selection and we suggest this as a complement to ongoing efforts to examine the function of ADPr-dependent signaling.

## ASSOCIATED CONTENT

### Supporting Information

Additional experimental details, materials, and methods; sequences and expected sizes of peptide substrates (Table S1), TLC-MALDI analysis of peptide modification by P15 (Figure S2), and non-enzymatic hydrolysis assay of P14p8 (Figure S3) (PDF)

The Supporting Information is available free of charge on the ACS Publications website.

## AUTHOR INFORMATION

### Author Contributions

The manuscript was written by Z.J. and I.C.-O. Experiments were designed by I.C.-O., Z.J., and S.R.W. Experiments were performed by Z.J., H.N., K.H., C.M., P.V., C.S., S.R.W., C.S., S.T., and A.F. and they analyzed the resulting data. All authors have given approval to the final version of the manuscript.

### Notes

The authors declare no competing financial interest.

## ACKNOWLEDGMENT

We thank K. Wheeler, P. Abbyad, A. Fuller, and M. Cohen for many helpful discussions regarding the manuscript and experimental design. We thank M. Cohen for supplying the *macro*D2 and TARG1 expression constructs. This work was funded by the National Institutes of Health (NIH 1R15GM139150) to I.C.-O.; Z.J., H.N., P.V., C.S. S.R.W., and S.T. were supported by internal funding from Santa Clara University.

---

Insert Table of Contents artwork here

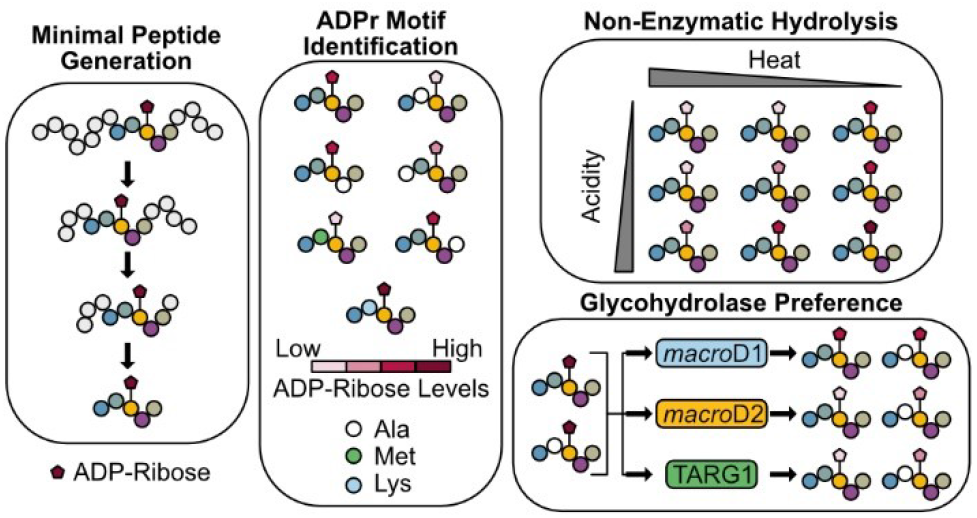

