## Supplemental Methods and Supplemental Figures for "Identification of a Novel PARP14 Site Motif and Glycohydrolase Specificity Using TLC-MALDI-TOF"

### Supporting Information

#### Protein and Peptide Preparation

Recombinant proteins were preceded by either a 6x His tag (P14, P15, or *macro*D1) or GST tag (*macro*D2 or TARG1) and were expressed and purified as previously described.[1] Peptides lacking a Y or W residue had a W added to the C-terminus to aid in concentration determination. The peptides were obtained from GenScript.

#### Chemoenzymatic Synthesis of ADPr—Peptide

P14 (8 µM) was incubated with 4 mM $NAD^+$ and either P14p6 or P14p8 (115 µM) for 2 hours at room temperature in a 420 µL reaction volume consisting of 50 mM HEPES, pH 7.5, 400 mM NaCl, 2.5% glycerol, and 0.4 mM β-mercaptoethanol. At 30 min an additional 11.8 µL of concentrated P14 (285 µM) was added. The reaction was diluted to 1 mL in SE buffer (10 mM potassium phosphate, pH 6.0 and 25 mM NaCl) and subjected to size exclusion chromatography at a rate of 0.75 mL/min using a Superdex Peptide 10/300 GL column (Cytiva) while collecting 250 µL fractions. ADPr—peptide containing fractions were pooled and brought up to 5 mL in IE buffer (10 mM BIS-Tris, pH 5.0 and 25 mM NaCl) and further purified using anion exchange chromatography with a HiTrap Q HP column (Cytiva) and exchanged into IE buffer + 1 M NaCl. ADPr—peptide fractions were desalted using a Sep-Pak C18 cartridge (Waters). Desalted peptides were diluted 1:1 with 100% acetonitrile, flash-frozen in liquid nitrogen, and lyophilized. Samples were stored at -80 °C immediately following drying to prevent ADPr hydrolysis.

#### TLC-MALDI Preparation and Sample Cleanup

The ultra-thin layer is prepared as previously described.[2] 2 µL of sample in MB is spotted directly on the thin-layer and dried. After drying, spots are washed twice with 5 µL of ice-cold 0.1% TFA for 1 minute. Calibrations were performed using Peptide Calibration Standard II (Bruker) per the manufacturer's instructions. MALDI-TOF experiments were performed using a MALDI-8020 instrument (Shimadzu).

#### MS Acquisition Parameters and Data Analysis

Collection of peptide spectra was performed using an automated firing method defined by a regular circle with a TV Raster (1500 µm diameter, 50 points, no offsets, 199 µm spacing) and random dithering (250 ms dwell time, 20 µm radius) with a mass range set to 700 – 2800 Da. 50 laser shots were fired for each profile at a frequency of 50 Hz and 2500 total shots were collected per sample. Post-acquisition baseline subtraction and smoothing were performed using MALDI Solutions (Shimadzu) with the following parameters: baseline filter width set between 15-30, Savitsky-Golay smoothing with a smoothing width between 5-30, and peak width set to 5. The peak delimiter method was set to threshold apex with a threshold of 0.01 mV and exclusion of peaks with intensities less than 0.01 mV. Following data acquisition, selected unmodified and ADPr-modified peaks were identified based on their expected mass and integrated to determine the area under the curve using MALDI Solutions and the resulting values were used to calculate relative levels of ADP-ribosylation. The center of each peak was recorded and spectra were rejected if any observed mass differed from the predicted mass by more than 1%. The spectral intensities were min-maxed normalized between 0.0 and 0.1 in Origin 2020 (OriginLab) and centered on the unmodified peak to aid comparisons between different peptides. Student's *t*-tests were performed with a two-tailed distribution to determine significance for any observed differences between experimental conditions in Excel (Microsoft). Linear fitting of non-enzymatic ADPr hydrolysis to a pseudo first-order rate expression was performed in Origin 2020.

| Peptide | Peptide Sequence | Unmodified m/z | +ADPr m/z | pI |
|---|---|---|---|---|
| P14p1 | SLNYKSTSSGHR**E**ISSPR | 2006.16 | 2546.16 | 10.49 |
| P14p2 | WHR**E**ISSPR | 1166.59 | 1706.59 | 10.95 |
| P14p3 | SLNYKSTSSGHR | 1336.43 | 1876.43 | 10.50 |
| P14p4 | RWSSGHR**E**ISS | 1301.38 | 1841.38 | 10.95 |
| P14p5 | SLNYKSTSSGHR**E**IS | 1664.81 | 2204.81 | 9.67 |
| P14p6 | HR**E**ISW | 826.91 | 1366.91 | 7.90 |
| P14p7 | AR**E**ISWH | 897.99 | 1437.99 | 7.90 |
| P14p8 | HA**E**ISWR | 897.99 | 1437.99 | 7.90 |
| P14p9 | HR**E**ASW | 784.83 | 1324.83 | 7.90 |
| P14p10 | HR**E**IAW | 810.91 | 1350.91 | 7.90 |
| P14p11 | HM**E**ISWR | 958.11 | 1498.11 | 7.90 |
| P14p12 | HK**E**ISW | 798.90 | 1338.90 | 7.88 |

**Table S1.** Peptides analyzed in the current study. Peptide sequence, unmodified *m/z*, expected +ADPr *m/z*, and isoelectric point (pI) are indicated.

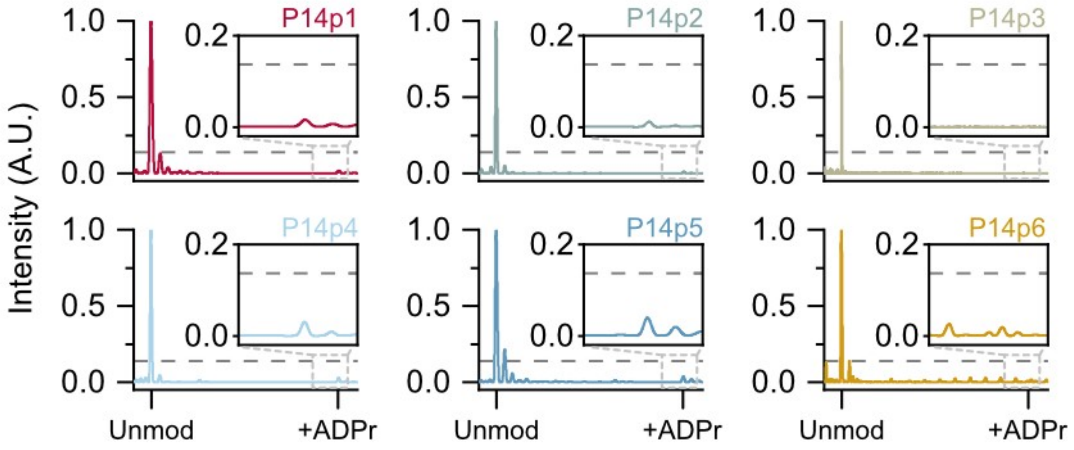


**Figure S2.** Minimal ADP-ribosylation of P14 selective peptides by P15. P15 and the indicated peptides were incubated in the presence of $NAD^+$ and subjected to TLC-MALDI to visualize the resulting increase in *m/z* due to ADPr (+541 Da). The dashed line represents the intensity observed for ADP-ribosylation of P14p6 by P14 and the inset highlights the +ADPr spectra.

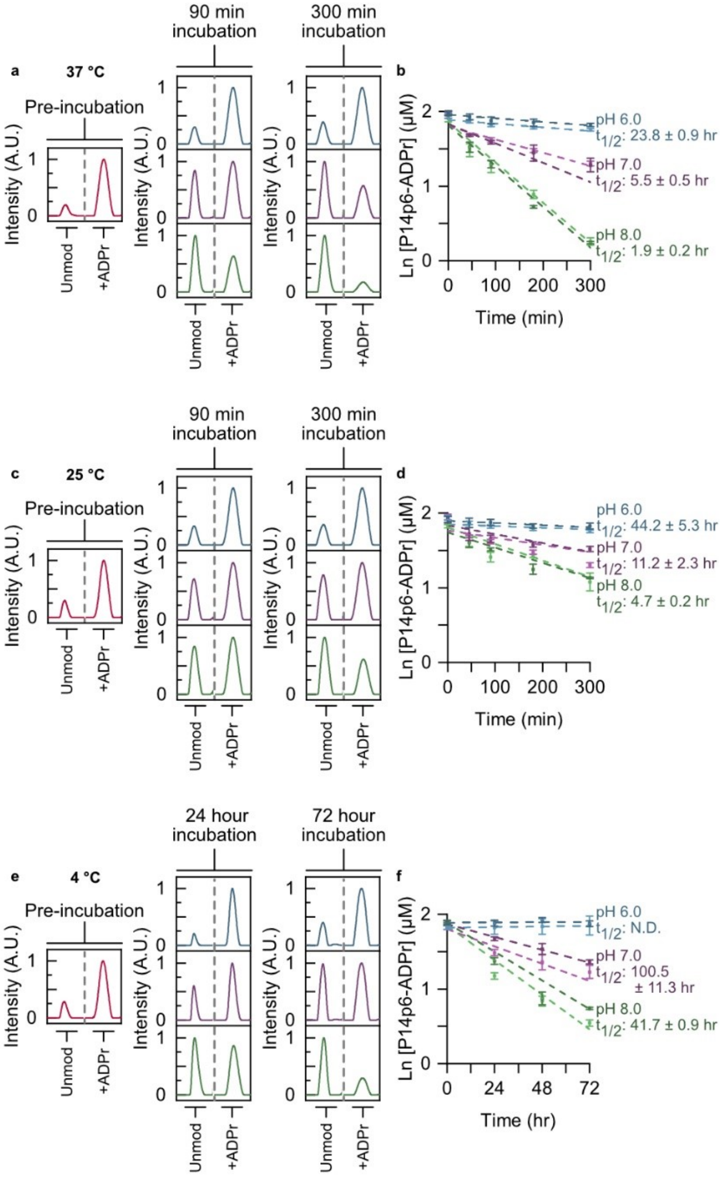


**Figure S3.** Non-enzymatic hydrolysis of ADPr is sequence independent. (a) Synthesized ADPr—P14p8 peptide was equilibrated at 37 °C and pH 6.0 (blue), pH 7.0 (purple), or pH 8.0 (green) and ADPr hydrolysis was monitored at the indicated times using TLC-MALDI. The unmodified and modified peaks are shown for comparison. (b) MS spectra were integrated to determine the relative levels of ADPr hydrolysis and fit to a pseudo first-order rate expression to determine the half-life of the ADPr modification (mean ± S.E.M., $n$ = 3). The fits obtained for P14p6 are shown for comparison (lighter coloring). (c) Experiments were performed as in (a) at 25 °C. (d) Determination of ADPr half-lives at 25 °C. (e) Experiments were performed as in (a) at 4 °C. (f) Determination of ADPr half-lives at 4 °C.
